# Predicting the Disposition of the Antimalarial Drug Artesunate and its Active Metabolite Dihydroartemisinin Using Physiologically-Based Pharmacokinetic Modeling

**DOI:** 10.1101/2020.10.28.360156

**Authors:** Ryan Arey, Brad Reisfeld

## Abstract

Artemisinin-based combination therapies (ACTs) have proven to be effective in helping to combat the global malaria epidemic. To optimally apply these drugs, information about their tissue-specific disposition is required, and one approach to predict these pharmacokinetic characteristics is physiologically-based pharmacokinetic (PBPK) modeling. In this study, a whole-body PBPK model was developed to simulate the time-dependent tissue concentrations of artesunate (AS) and its active metabolite, dihydroartemisinin (DHA). The model was developed for both rats and humans and incorporated drug metabolism of the parent compound and major metabolite. Model calibration was conducted using data from the literature in a Bayesian framework, and model verification was assessed using separate sets of data. Results showed good agreement between model predictions and the validation data, demonstrating the capability of the model in predicting the blood, plasma, and tissue pharmacokinetics of AS and DHA. It is expected that such a tool will be useful in characterizing the disposition of these chemicals and ultimately improve dosing regimens by enabling a quantitative assessment of the tissue-specific drug levels critical in the evaluation of efficacy and toxicity.

## Introduction

Malaria is a global health epidemic resulting in the deaths of nearly half a million people per year (**1**). The World Health Organization (WHO) recommends artemisinin-based combination therapies (ACTs) as a first-line treatment against uncomplicated *Plasmodium falciparum* malaria. In countries where malaria is endemic, treatment policies have been progressively updated with the implementation of ACTs in lieu of monotherapies such as chloroquine, amodiaquine, and sulfadoxine-pyrimethamine, leading to a substantial reduction in global morbidity and mortality (**1**). Artemisinin and semisynthetic derivatives such as, artesunate (AS), artemether (AM), and dihydroartemisinin (DHA), are short-acting antimalarial agents that kill the parasites more rapidly than conventional antimalarial drugs and are active against asexual and some sexual forms of the parasite (**2**).

Artemisinin is a sesquiterpene lactone endoperoxide. Though the mechanism of action is not completely understood, it has been shown that antimalarial activity results from cleavage of the peroxide bridge in the presence of ferrous iron (Fe^+2^), producing reactive oxygen species, which are thought to mediate the cytotoxic effect (**3**). The chemical compounds resulting from metabolism of artemisinins generally fall into one of two categories: (i) hydroxylated compounds with the peroxide bridge intact, which are biologically active, and (ii) deoxy compounds with a reduced peroxide bridge, forming biologically inactive compounds. All biologically active metabolites undergo further metabolism via glucuronidation and/or other conjugation, and eventual excretion through the urine and feces (**4**). Following intravenous (IV) administration of the semisynthetic AS, the parent compound is rapidly converted to its active metabolite, DHA (**2**). Both compounds are available through various routes of administration, but IV AS is the WHO’s recommended treatment option for severe malaria because it’s the only artemisinin derivative with sufficient water solubility (**5**). Unfortunately, despite the overall effectiveness of ACTs, concerns have been raised about their potential for toxicity in numerous vital organs, including the brain, heart, and kidneys, as well as a potential for embryotoxicity, genotoxicity, and hemato-immunotoxicity (**13**). AS has demonstrated cytotoxicity in mammalian cells via apoptosis as the main route of cell death, indicating a potential for adverse side effects (**13**).

Characterizing drug ADME (absorption, distribution, metabolism, and excretion) is critical in developing appropriate dosing strategies to assure safety and efficacy. As a result, both experimental and mathematical modeling studies have been conducted to elucidate the pharmacokinetic characteristics of both AS and DHA. Though most of the experimental pharmacokinetic studies for these compounds have focused on acquiring drug concentrations in plasma over time (**6–12**), a few investigators have collected data in other tissues (**9, 10, 12**). These latter studies have all been conducted inrats utilizing radiolabeled doses of the drug and determining concentration with respect to time through measurement of total radioactivity (TR). Complementing this research has been the development of non-compartmental or simple compartmental models based on the acquired data with the intent of estimating various PK metrics of interest, such as C_max_ and AUC. A more mechanistically-detailed model was created by Gordi et al. (**14**) who included in their representation the effects of autoinduction in the liver and were able to accurately predict the PK behavior of artemisinin in healthy human subjects. More recently, Olafuyi and coworkers (**15**) utilized the Advanced Dissolution, Absorption, and Metabolism (ADAM) model (**16**) to predict the pharmacokinetics of artemether-lumefantrine combination therapies in both adult and pediatric populations, including potential drug-drug interactions of lumefantrine and the tuberculosis chemotherapy, rifampicin.

Overall, though informative experiments and modeling studies have been conducted to characterize the disposition of AS, there remain significant questions. In particular, from the perspective of both efficacy and safety, what are the concentrations of AS and its active metabolites over time in tissues relevant to humans? To help address this key question, the aims of the current study were to develop a whole-body physiologically-based pharmacokinetic (PBPK) model capable of describing the kinetics involved in the ADME of AS and its active metabolite, DHA. To this end, the overall strategy was to first developed and validate a model for rats, to extrapolate the model to humans, and conduct verification based on available data.

## Materials and Methods

### Approach

To achieve the study aims, two generic whole-body PBPK models were developed, parameterized, and validated: (i) a rat-specific PBPK model (R-PBPK) and (ii) a human-specific PBPK model (H-PBPK). Both models shared the same compartmental structure and governing equations with the only difference being values of parameters related to the anatomy, physiology, and metabolism of drugs by each biological species. The models were parameterized within a Bayesian framework for both species by utilizing sets of training data mined from the literature. Models were validated using separate data sets. Here, the term *validation* refers to confirmation of the plausibility of the proposed model in representing the underlying real system, as described by Tomlin and Axelrod (2007). In this paper, the terms *validation* and *verification* will be used interchangeably to describe the process of determining if the model, as constructed accurately, represents the underlying real system being modeled by comparing the simulation output with experimental data from the real system that were not used in the parameterization process.

### Training and Validation data

A summary of the data used in this study are shown in **Table 1**. In more specific terms, pharmacokinetic data for calibration of the R-PBPK model were obtained from various papers (**9–10**) that provided (i) drug concentrations in tissues relevant to the efficacy or toxicity of the drug in the human body and (ii) information on the fraction of drug bound to plasma proteins for both AS and DHA. These data were compiled to generate a training set for parameterization of the rat model. For training of the human model, a thorough literature search was conducted to identify studies (**6, 18**) that provided plasma drug concentrations of AS or DHA with emphasis placed on those studies that utilized an IV route of administration. These data comprised the training set for the calibration of the human model. For model validation, additional data were utilized (**7, 8, 11, 12**) with the same selection criteria as above and set aside for testing and verification of model predictions.

**TABLE 1.**
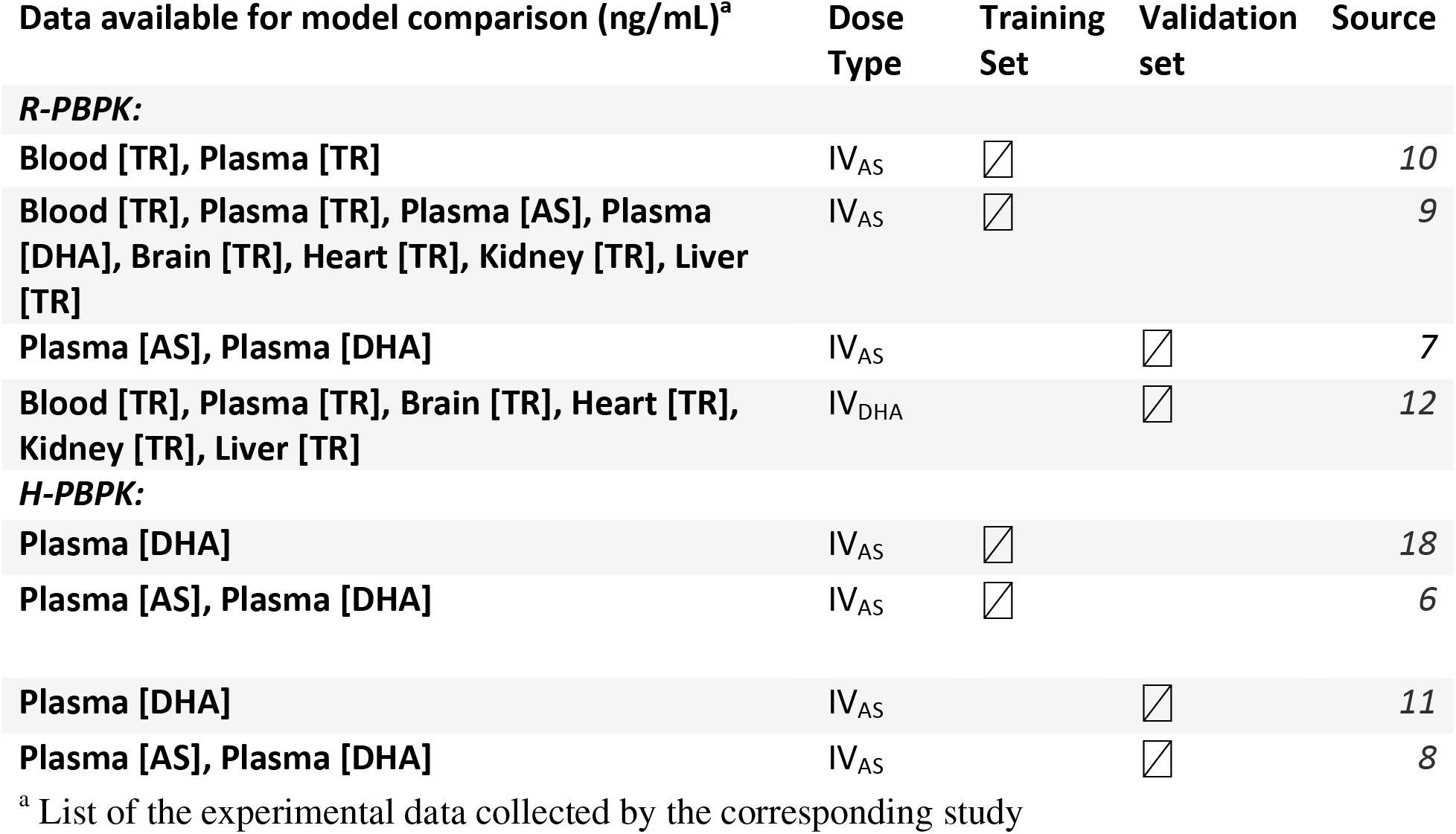
Data used for training (calibration) and validation of both models

| Data available for model comparison (ng/mL)<sup>a</sup> | Dose Type | Training Set | Validation set | Source |
| --- | --- | --- | --- | --- |
| ***R-PBPK:*** |  |  |  |  |
| Blood [TR], Plasma [TR] | IV<sub>AS</sub> | ☒ |  | 10 |
| Blood [TR], Plasma [TR], Plasma [AS], Plasma [DHA], Brain [TR], Heart [TR], Kidney [TR], Liver [TR] | IV<sub>AS</sub> | ☒ |  | 9 |
| Plasma [AS], Plasma [DHA] | IV<sub>AS</sub> |  | ☒ | 7 |
| Blood [TR], Plasma [TR], Brain [TR], Heart [TR], Kidney [TR], Liver [TR] | IV<sub>DHA</sub> |  | ☒ | 12 |
| ***H-PBPK:*** |  |  |  |  |
| Plasma [DHA] | IV<sub>AS</sub> | ☒ |  | 18 |
| Plasma [AS], Plasma [DHA] | IV<sub>AS</sub> | ☒ |  | 6 |
| Plasma [DHA] | IV<sub>AS</sub> |  | ☒ | 11 |
| Plasma [AS], Plasma [DHA] | IV<sub>AS</sub> |  | ☒ | 8 |

<sup>a</sup> List of the experimental data collected by the corresponding study

### PBPK model

As shown in **Figure 1,** the structural model comprised compartments for arterial and venous blood, lung, heart, brain, muscle, fat, bone, spleen and pancreas (single compartment), liver, kidney, and the gut, as well as a compartment for the rest of the body containing tissues not accounted for in other compartments. All compartments were assumed to be perfusion-limited, and as noted earlier, this structure was identical for both the rat and human models. Components related to the metabolism of AS are described below. Enterohepatic recirculation (EHR) of drugs was modeled by biliary excretion of conjugated metabolites (**19**) to a sub-compartment of the gut, representing the lumen of the small intestine. The governing equations embodying these biological features form a system of mass-balanced ordinary differential equations (ODE’s), dependent on model parameters and time. These equations dictate the rate of change in the amount of drug in each compartment. See the appendix for a description of the model equations.

**FIGURE 1.**
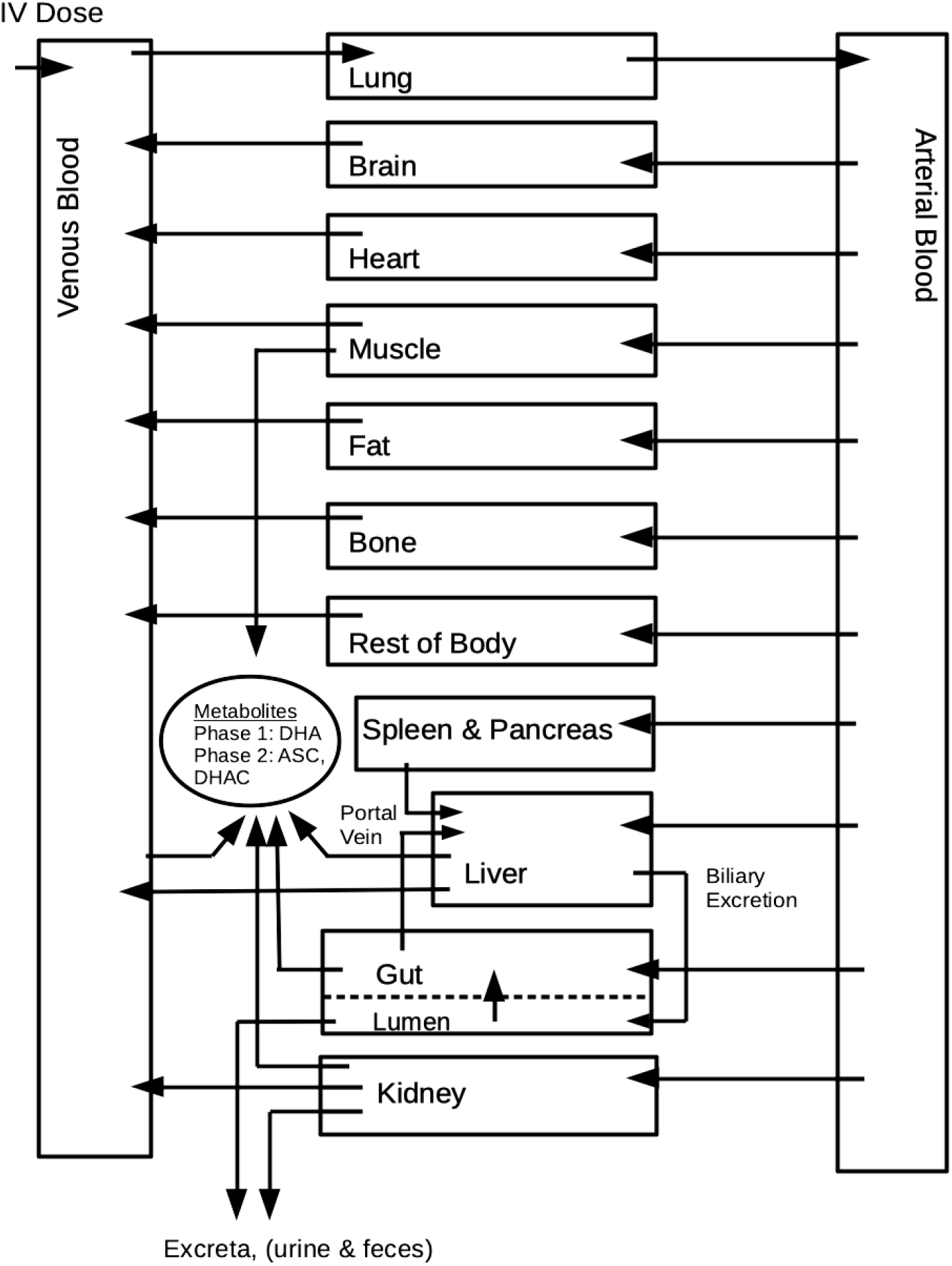
Schematic detailing the generic whole-body structure of the PBPK model.

### Drug metabolism

The drug AS is subjected to both phase 1 and phase 2 reactions, where the phase 1 reactions yield DHA and the phase 2 reactions yield a conjugated product of AS or DHA (**9**). In this work, we refer to the conjugated product of AS as *AS-C* and the conjugated product of DHA as *DHA-C*.

The metabolism of AS and DHA was assumed to occur in the liver and a variety of extrahepatic tissues, namely blood, muscle, gut, and kidneys. Specifically, it was assumed that AS was metabolized immediately upon entering the blood, being hydrolyzed to DHA in the venous compartment, as well as being metabolized in the liver compartment, as was observed in the literature (**20–24**). Drug conjugation of both active chemical species (AS and DHA) was assumed to occur in muscle, gut, kidney, and liver tissues. This assumption was supported by findings from a study by Illett et al. (2002), which demonstrated that conjugation of DHA occurs via specific isoforms of uridine diphosphate glucuronosyltransferases (UGTs), namely UGT1A9 and UGT2B7, to form DHA-glucuronide (**24**), along with data supporting that these specific UGT isoforms are present in muscle, gut, kidney and liver tissue (**25**). We further assumed that AS was conjugated in the same tissues as DHA, and that the conjugated product of both species (AS-C and DHA-C) represent a nonspecific, lumped term accounting for all drug conjugates of that particular chemical compound. An overall schematic of the assumed metabolic scheme is shown in **Figure 2**.

**FIGURE 2.**
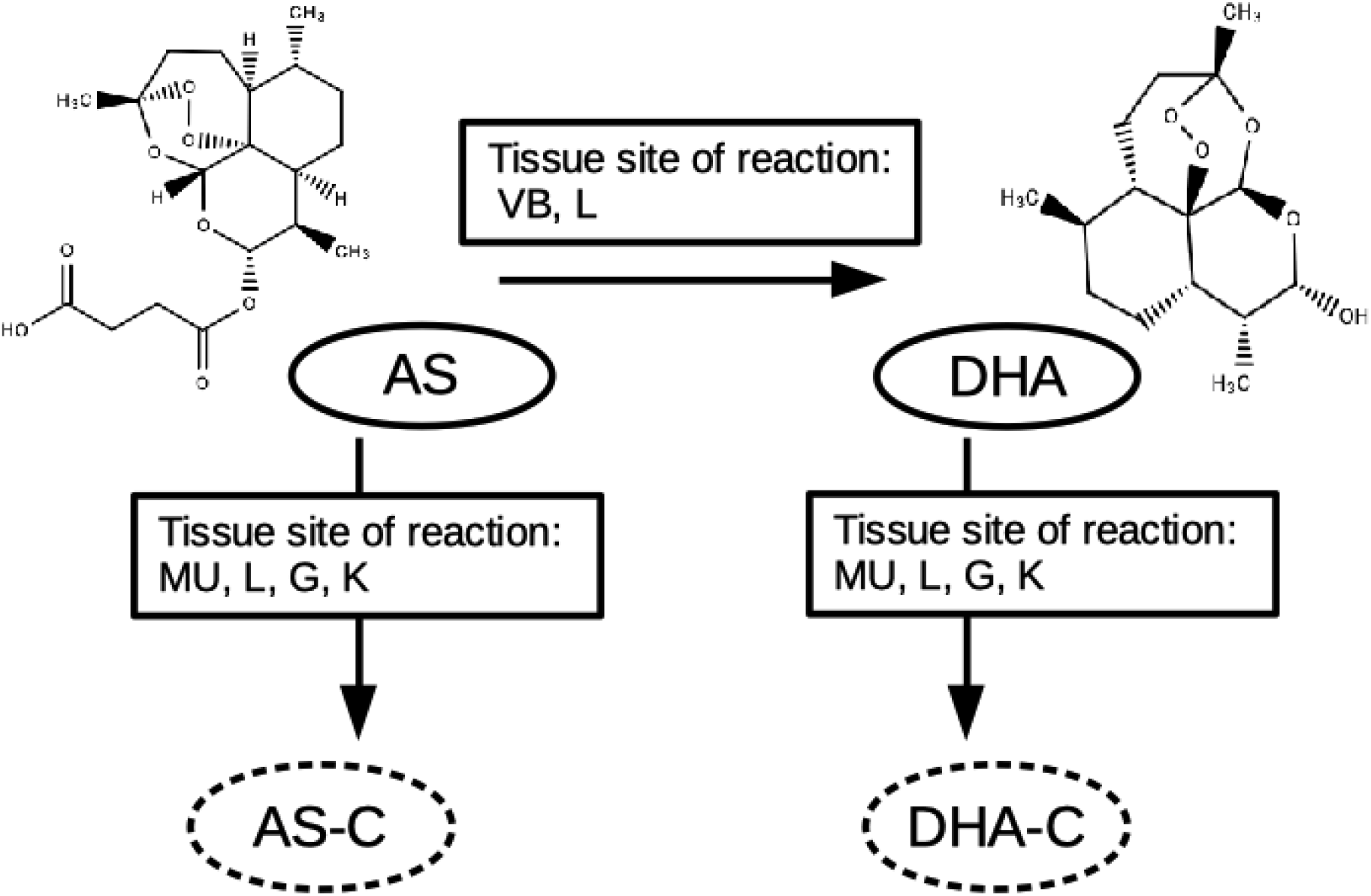
Diagram of the metabolic pathways of AS and DHA listed with the corresponding hypothesized tissue site of metabolism, where VB, MU, L, G, K represent venous blood, muscle, liver, gut, and kidney tissues, respectively. Biologically active compounds are encircled by a solid line while biologically inactive compounds are encircled with a dotted line.

Based on results from the literature (**24, 26, 27**), all equations pertaining to the metabolism of AS and DHA were assumed to follow Michaelis-Menten (M-M) reaction kinetics, modeled as

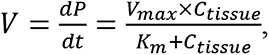

where the rate of product formation 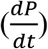 relies upon the maximum velocity of the reaction rate (V_max_), the Michaelis constant (K_m_), and drug concentration at the tissue site of metabolism (C_tissue_).

### Specification of parameter values

Anatomical and physiological parameters were obtained from Brown et al. (1997) and Delp et al. (1991). Organ/tissue volumes were scaled linearly with body weight while blood flow rates were allometrically scaled with body weight to the 0.75 power (**28–30**). Tissue density was assumed equal to that of water (~1 g/mL). Fraction bound parameters and clearance parameters were taken from the literature, where clearance via renal and biliary excretion was scaled by apportioning a fraction of total blood clearance to the kidneys, with the remaining fraction being that for the liver (**9**). M-M parameters for the metabolism of AS and DHA in the liver compartment were taken from *in vitro* experimental results (**24, 26, 27**) derived using human liver microsomes and recombinant UGTs. These values were then scaled to *in vivo* conditions for model simulation using information from other studies (**31, 32**). Metabolic rates for blood, muscle, gut, and kidney compartments were determined from a nonlinear least square fit of model-simulated data following M-M reaction kinetics in each extrahepatic tissue. Moreover, metabolism in these tissues was assumed to be proportional to the known metabolic rates of each compound in the liver. This assumption was incorporated by estimating coefficients assigned to the metabolic equations in each of the extrahepatic tissues (see **Table 2**). First-order rate constants for absorption and excretion of drugs from the gut lumen were calculated from information found in the literature (**20, 31**). Values for the tissue/plasma partition coefficients of AS and DHA were computed using the httk (v2.0.1) package for the statistical software R (v3.6.1) (**33**), while partition coefficients for the lumped, conjugated terms (AS-C and DHA-C) were estimated from tissue concentration-time data (**9**). Specifically, the conjugated partition coefficients were computed as *P* = *C*_*tissue*_/*C*_*plasma*_, where *P* = partition coefficient, *C*_*tissue*_ = drug concentration (TR) in tissue, and *C*_*plasma*_ = drug concentration (TR) in plasma. The mean values were computed using time points during the elimination phase at which equilibrium in drug concentration was assumed between the tissues and venous blood. For the conjugated terms, partition coefficients of the ‘REST’ compartment (rest of the body) were set equal to the weighted mean of all other partition coefficients. All model parameters are listed with their values and corresponding sources in **Table 2**.

**TABLE 2.** All model parameters listed with their corresponding values pulled from the literature^*a*^

| Parameter description with units | Parameter symbol | AS values | DHA values | Source |
| --- | --- | --- | --- | --- |
| Molecular weight (mg/mmol) | $MW$ | 384.4 | 284.9 | 7 |
| Logarithm of octanol:water partitioning | $\log P$ | 2.05 | 2.19 | 33, 50 |
| Dissociation constant (acid,base) | $pK_a$ | 3.77, -4.2 | 12.11, -4.1 | 33, 51 |
| Plasma binding components | AAG | α1-acid-glycoprotein | α1-acid-glycoprotein | 2 |
| Anatomical and physiological parameters: |  | R-PBPK values | H-PBPK values |  |
| Body Weight (kg) | $BW$ | 0.25 | 70 | 28 |
| Cardiac Output (L/min) | $QCC$ | 0.235 | 0.215 | 29 |
| Fractional volumes (fraction of BW) |  |  |  |  |
| Lung | $FV_{Lu}$ | 0.005 | 0.008 | 28 |
| Brain | $FV_{BR}$ | 0.006 | 0.02 | 28 |
| Heart | $FV_H$ | 0.003 | 0.005 | 28 |
| Muscle | $FV_{MU}$ | 0.404 | 0.4 | 28 |
| Fat | $FV_F$ | 0.07 | 0.214 | 28 |
| Bone | $FV_B$ | 0.073 | 0.143 | 28 |
| Pancreas and Spleen | $FV_{PS}$ | 0.0052 | 0.004 | 28 |
| Liver | $FV_L$ | 0.034 | 0.026 | 28 |
| Gut | $FV_G$ | 0.077 | 0.031 | 28 |
| Kidney | $FV_K$ | 0.007 | 0.004 | 28 |
| Total Blood | $FV_{BL}$ | 0.074 | 0.079 | 28 |
| Venous blood | $FV_{VB}$ | 0.65*FV_BL | 0.65*FV_BL | 30 |
| Arterial blood | $FV_{AB}$ | 0.35*FV_BL | 0.35*FV_BL | 30 |
| Rest of the body | $FV_{REST}$ | 1 - (all other tissues) | 1 - (all other tissues) | * |
| Fractional flow rates (fraction of Cardiac Output) |  |  |  |  |
| Brain | $FQ_{BR}$ | 0.02 | 0.114 | 28 |
| Heart | $FQ_H$ | 0.049 | 0.04 | 28 |
| Muscle | $FQ_{MU}$ | 0.278 | 0.191 | 28 |
| Fat | $FQ_F$ | 0.07 | 0.052 | 28 |
| Bone | $FQ_B$ | 0.122 | 0.042 | 28 |
| Hepatic artery | $FQ_{HA}$ | 0.024 | 0.046 | 28 |
| Portal vein | $FQ_{PV}$ | 0.151 | 0.181 | 28 |
| Gut | $FQ_G$ | 0.14 | 0.14 | 29 |
| Kidney | $FQ_K$ | 0.141 | 0.175 | 28 |
| Rest of the body | $FQ_{REST}$ | 1 - (all other tissues) | 1 - (all other tissues) | * |
| Fraction bound in plasma (dimensionless) |  |  |  |  |
| AS: | $fb_{AS}$ | 0.82 | 0.75 | 9 |
| DHA: | $fb_{DHA}$ | 0.82 | 0.75 | 9 |
| Fraction unbound in plasma (dimensionless) |  |  |  |  |
| AS: | $fub_{AS}$ | $1 - fb_{AS}$ | $1 - fb_{AS}$ | 9 |
| DHA: | $fub_{DHA}$ | $1 - fb_{DHA}$ | $1 - fb_{DHA}$ | 9 |
| Total blood clearance (mL/hr/kg) | $CLC_{TB}$ | 0.0815 | 0.0815 | 9 |
| Fractional renal clearance (dimensionless) | $f_r$ | 0.6 | 0.6 | 9 |
| Renal clearance (mL/hr/kg) | $CLC_{renal}$ | $f_r * CLC_{TB}$ | $f_r * CLC_{TB}$ | 9 |
| Biliary clearance (mL/hr/kg) | $CLC_{bile}$ | $(1 - f_r) * CLC_{TB}$ | $(1 - f_r) * CLC_{TB}$ | 9 |
| Michaelis-Menten metabolic parameters |  |  |  |  |
| AS: |  |  |  |  |
| (mg/hr/gram of Liver) | $Vmax_{AS2B6}$ | 0.1782 | 0.1148 | 27 |
| (mg/L) | $Km_{AS2b6}$ | 0.75 | 0.75 | 27 |
| "" | $Vmax_{AS3A4}$ | 0.072 | 0.0464 | 26 |
| "" | $Km_{AS3A4}$ | 42.7 | 42.7 | 26 |
| "" | $Vmax_{AS3A5}$ | 0.133 | 0.0857 | 26 |
| "" | $Km_{AS3A5}$ | 79.1 | 79.1 | 26 |
| DHA: |  |  |  |  |
| (mg/hr/gram of Liver) | $Vmax_{DHA1A9}$ | 0.0068 | 0.0044 | 24 |
| (mg/L) | $Km_{DHA1A9}$ | 9.1 | 9.1 | 24 |
| "" | $Vmax_{DHA2B7}$ | 0.0084 | 0.0054 | 24 |
| "" | $Km_{DHA2B7}$ | 125 | 125 | 24 |
| "" | $Vmax_{DHA}$ | 0.1359 | 0.08758 | 24 |
| "" | $Km_{DHA}$ | 25.59 | 25.59 | 24 |
| **Estimated metabolic rates in extrahepatic tissues** |  |  |  |  |
| (mg/hr) | $V_{max_{ASeff}}$ | 1.6 | 1.6 | * |
| (mg/L) | $K_{m_{ASeff}}$ | 0.78 | 0.78 | * |
| "" | $V_{max_{DHAeff}}$ | 1.2 | 1.2 | * |
| "" | $K_{m_{DHAeff}}$ | 26 | 26 | * |
| **Coefficients of unknown metabolic rates:** |  |  |  |  |
| AS metabolism (venous blood) | $C_{m1}$ | 2 | 150 | * |
| AS conjugation (muscle, liver, kidney, gut) | $C_{m2}$ | 30 | 10 | * |
| DHA conjugation (muscle, liver, kidney, gut) | $C_{m3}$ | 1 | 200 | * |
| Apparent permeability coefficient (cm/s) | $P_{app}$ | 4.00E-06 | 4.00E-06 | 20 |
| radius of small intestine (cm) | $radius$ | 0.18 | 1.0 | 31 |
| Gut lumen to Gut absorption rate (1/hr) | $K_{GLG}$ | (Papp/radius)*<br>60<sup>2</sup> | (Papp/radius)*<br>60<sup>2</sup> | 20, 31 |
| Gut excretion rate (1/hr) | $K_{Gut}$ | 2.1 | 2.1 | 31 |
| **Partition Coefficients: (dimensionless)** |  |  |  |  |
| AS: | $PLU_{AS}$ | 2.5 | 2.5 | 33 |
| | $PBR_{AS}$ | 1.5 | 1.5 | 33 |
| | $PF_{AS}$ | 0.65 | 0.65 | 33 |
| | $PMU_{AS}$ | 1.9 | 1.9 | 33 |
| | $PB_{AS}$ | 1.3 | 1.3 | 33 |
| | $PH_{AS}$ | 3.2 | 3.2 | 33 |
| | $PPS_{AS}$ | 2.7 | 2.7 | 33 |
| | $PK_{AS}$ | 9.7 | 9.7 | 33 |
| | $PL_{AS}$ | 11 | 11 | 33 |
| | $PG_{AS}$ | 3.3 | 3.3 | 33 |
| | $PREST_{AS}$ | 3.6 | 3.6 | 33 |
| **Represents blood-to-plasma ratio** | $PRBC_{AS}$ | 1.4 | 1.4 | 33 |
| DHA: | $PLU_{DHA}$ | 9.7 | 9.7 | 33 |
| | $PBR_{DHA}$ | 6.1 | 6.1 | 33 |
| | $PF_{DHA}$ | 14 | 14 | 33 |
| | $PMU_{DHA}$ | 3.9 | 3.9 | 33 |
| | $PB_{DHA}$ | 4 | 4 | 33 |
| | $PH_{DHA}$ | 6.2 | 6.2 | 33 |
| | $PPS_{DHA}$ | 5.2 | 5.2 | 33 |
| | $PK_{DHA}$ | 16 | 16 | 33 |
| | $PL_{DHA}$ | 18 | 18 | 33 |
| | $PG_{DHA}$ | 11 | 11 | 33 |
| | $PREST_{DHA}$ | 9.1 | 9.1 | 33 |
| **Represents blood-to-plasma ratio** | $PRBC_{DHA}$ | 1.9 | 1.9 | 33 |
| **AS-C & DHA-C:** | $PLU_{AS-C, DHA-C}$ | 4 | 4 | 9 |
| | $PBR_{AS-C, DHA-C}$ | 1.6 | 1.6 | 9 |
| | $PF_{AS-C, DHA-C}$ | 0.7 | 0.7 | 9 |
| | $PMU_{AS-C, DHA-C}$ | 1.5 | 1.5 | 9 |
| | $PB_{AS-C, DHA-C}$ | 2.1 | 2.1 | 9 |
| | $PH_{DAS-C, DHA-C}$ | 6.1 | 6.1 | 9 |
| | $PPS_{AS-C, DHA-C}$ | 24.5 | 24.5 | 9 |
| | $PK_{AS-C, DHA-C}$ | 18.6 | 18.6 | 9 |
| | $PL_{AS-C, DHA-C}$ | 9.3 | 9.3 | 9 |
| | $PG_{AS-C, DHA-C}$ | 84.5 | 84.5 | 9 |
| | $PREST_{AS-C, DHA-C}$ | 8.2 | 8.2 | 9 |
| **Represents blood-to-plasma ratio** | $PRBC_{AS-C, DHA-C}$ | 1.9 | 1.9 | 33 |

<sup>a</sup> Values displayed here represent the values of each parameter before scaling to *in vivo* conditions for simulation.
\* Estimated from the present study

### Sensitivity analysis

A local sensitivity analysis was performed utilizing a method reported in previous PBPK research (**34**). This task was accomplished by using a Monte Carlo (MC) approach, wherein each model parameter was uniformly distributed between a lower bound (half the mean parameter value) and an upper bound (double the mean parameter value) for 15,000 MC draws. All model parameters were sampled simultaneously to produce a distribution of model outputs. Mean PK parameters, namely maximum concentration (C_max_), time to maximum concentration (T_max_), and area under the concentration-time curve (AUC) were then calculated from the distributed time course concentration data in the MC output. Pearson correlation coefficients were calculated between model parameters and mean PK parameters, while those returning an absolute value of ≥ 0.2 were considered to be influential.

### Simulation methodology and computing platform

Parameter estimation was carried out in a Bayesian hierarchical context using Markov Chain Monte Carlo (MCMC) simulations applying the Metropolis-within-Gibbs algorithm. Calibration of the R-PBPK model was conducted first from prior distributions of the influential parameters constructed around their respective source values over the course of 200,000 iterations. Convergence was assessed by the potential scale reduction factor (**35**) by producing three separate chains from a two-level, hierarchical MCMC simulation. The model equations were then computed by sampling the influential parameters from their corresponding posterior distributions at the last 10,000 iterations, along with the distribution of other relevant partition coefficients for comparison to the validation data pulled from the literature. For calibration of the H-PBPK model, all parameters associated with the metabolism or excretion of drugs were estimated in the same Bayesian context over the same number of iterations. Influential parameters were held at constant values set equal to the mean of the posterior resulting from the calibration step of the R-PBPK model. The estimated human parameters were then sampled from their corresponding posterior distributions at the last 10,000 iterations, resulting from the calibration step of the H-PBPK model, for comparison to the validation dataset extracted from the literature (see **Table 3**).

**TABLE 3.**
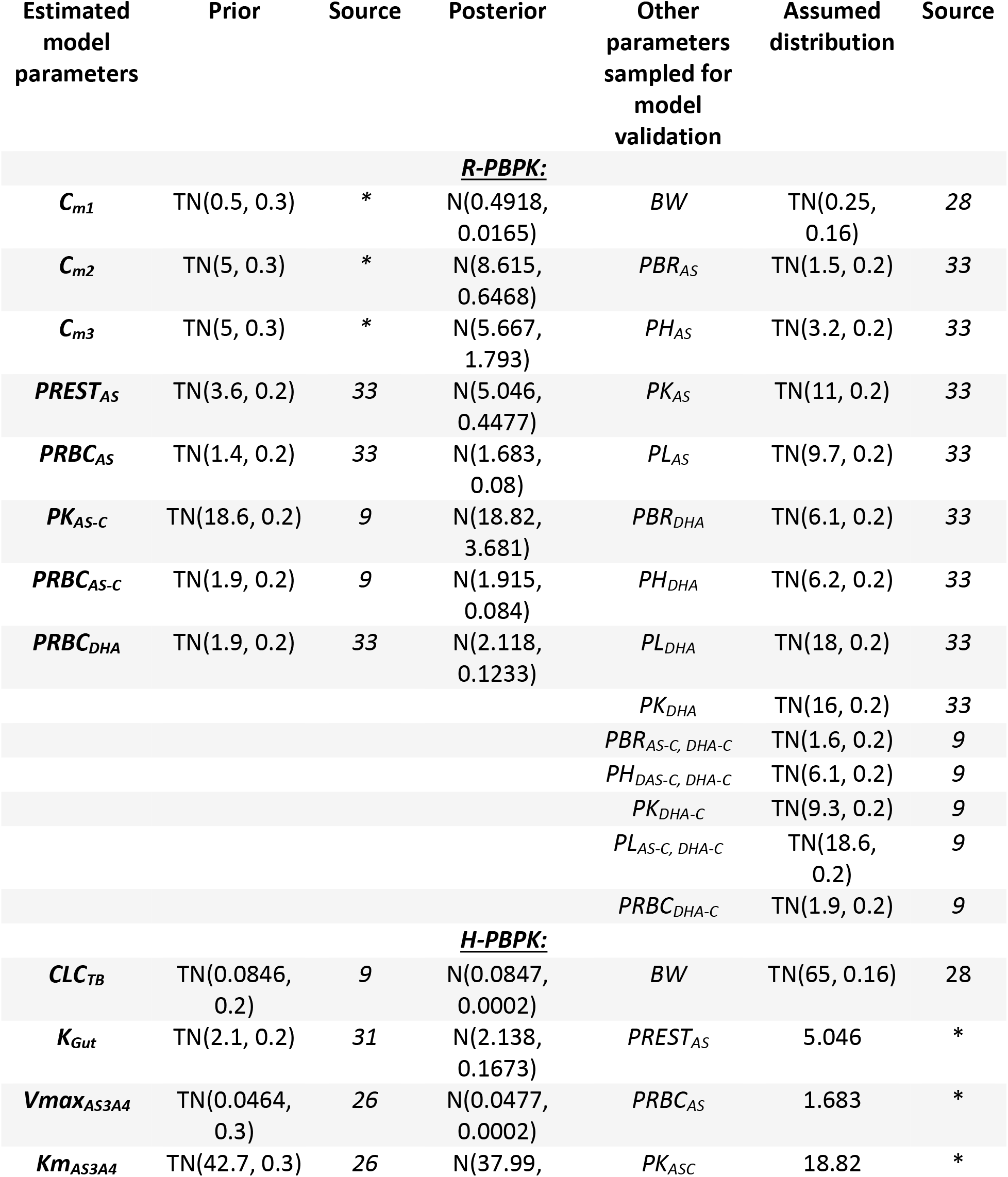
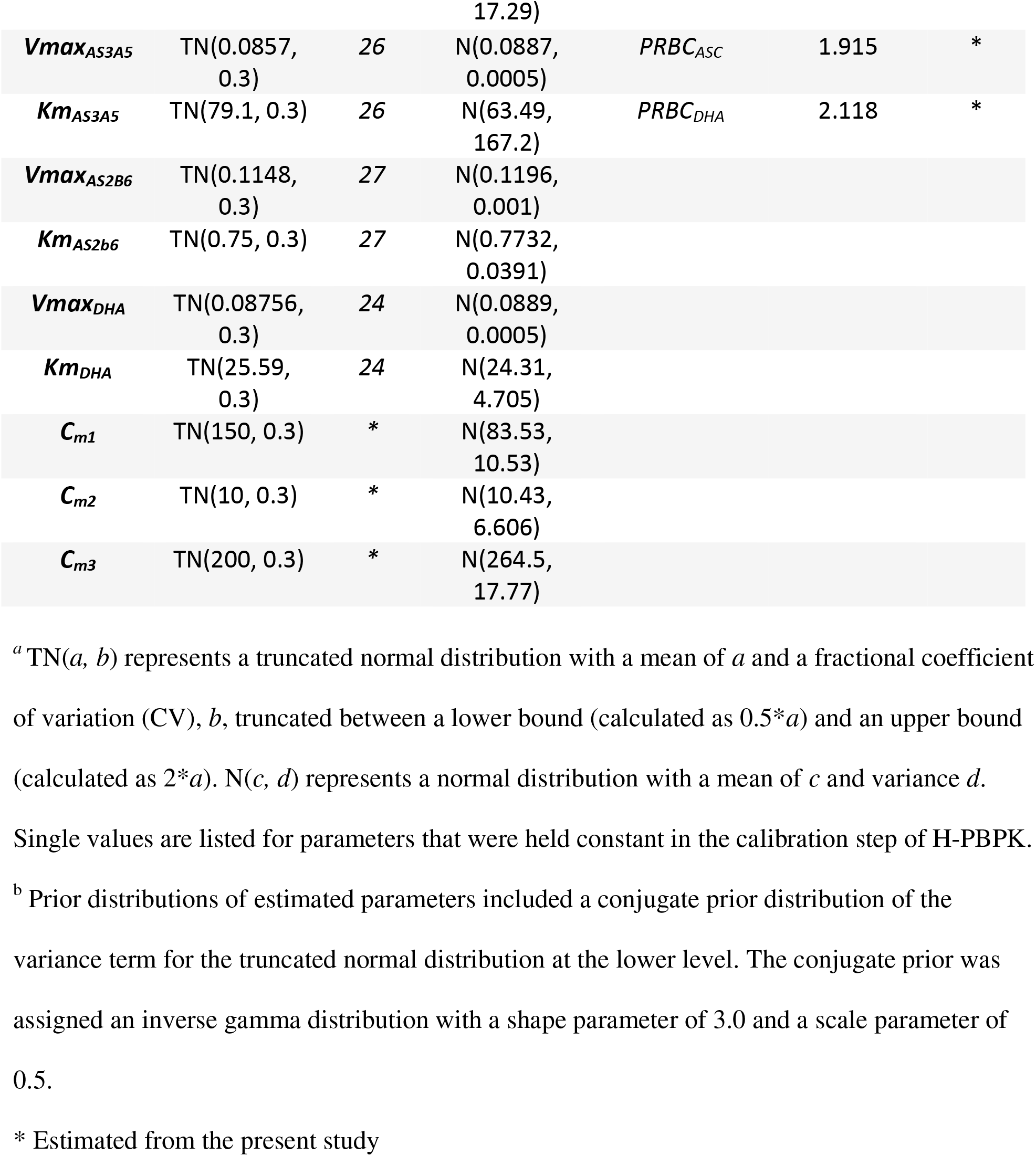
Probability distributions of model parameters for the purposes of model calibration (priors)^a,b^ and model validation (posteriors and the assumed distributions of other relevant parameters)

Data from the literature were digitized using Engauge Digitizer (v12.1) (**36**). Simulations of the PBPK governing equations, including MC and MCMC simulations, were conducted in MCSim (v6.1.0) (**37**). Data-processing, analysis, and visualization of data were carried out in Python (v3.6.8) (**38**) utilizing the pandas (**39),** numpy (**40**), scipy (**41**), and matplotlib (**42**) packages, the statistical software R (v3.6.1) (**43**), utilizing the httk (**33**), coda (**44**), and ggplot2 packages, and finally Microsoft Excel (v16.34) (**46**). All computations were performed on general compute nodes running the Red Hat Enterprise Linux 7 operating system, each containing Intel Xeon E5-2680 v3 processors at 2.5GHz with 4.84 GB of RAM per core, with a total of 24 cores per node.

## Results

### Model parameters

Using the procedure outlined above, influential model parameters were estimated via Bayesian inference from their respective prior distributions. All computed prior and posterior distributions are listed in **Table 3**. For the case of the R-PBPK model, the posterior distributions correspond to MCMC simulations in which the potential scale reduction factor (PSRF) was computed as an R value of 1.0 for each of the estimated model parameters, suggesting that each of the posteriors had converged to a stationary distribution (**47**). With the H-PBPK model, convergence was observed for most of the estimated parameters by the PSRF obtained as an R value between 1.0 and 1.05 with the exception of four parameters, namely two of the Michaelis constants for the metabolism of AS in the liver (*Km*_*AS3A4*_ and *Km*_*AS3A5*_), the scaling coefficient of AS metabolism in the blood (*Cm*_*1*_), and the scaling coefficient of DHA conjugation in the extrahepatic tissues (*Cm*_*3*_). The nonconverging parameters returned R values corresponding to the PSRF of 1.5, 1.4, 4.41, and 9.76, respectively, indicating that a stationary distribution had not been reached. The reasons behind this lack of convergence for certain parameter distributions are unclear at present.

### Testing and validation of the R-PBPK model

The R-PBPK model was evaluated by sampling (i) the influential parameters from their corresponding posterior distributions and (ii) relevant partition coefficients from distributions of their computed mean values (**Table 3**). The simulation results produced a range of R-PBPK model outputs for comparison to the *in vivo* concentration-time data selected for model validation (**7, 12**). Data selected for model comparison included AS and DHA concentrations in plasma, as well as TR concentrations in blood, plasma, brain, heart, liver, and kidney tissues. This comparison is illustrated in **Figures 3** and **4**, which shows the validation data (points) co-plotted with the mean concentration-time curve (solid line) and 95% prediction intervals (greyed area) of the R-PBPK model. For comparison of these data, IV doses of AS and DHA were simulated as a short bolus over a duration less than 1 minute.

**FIGURE 3.**
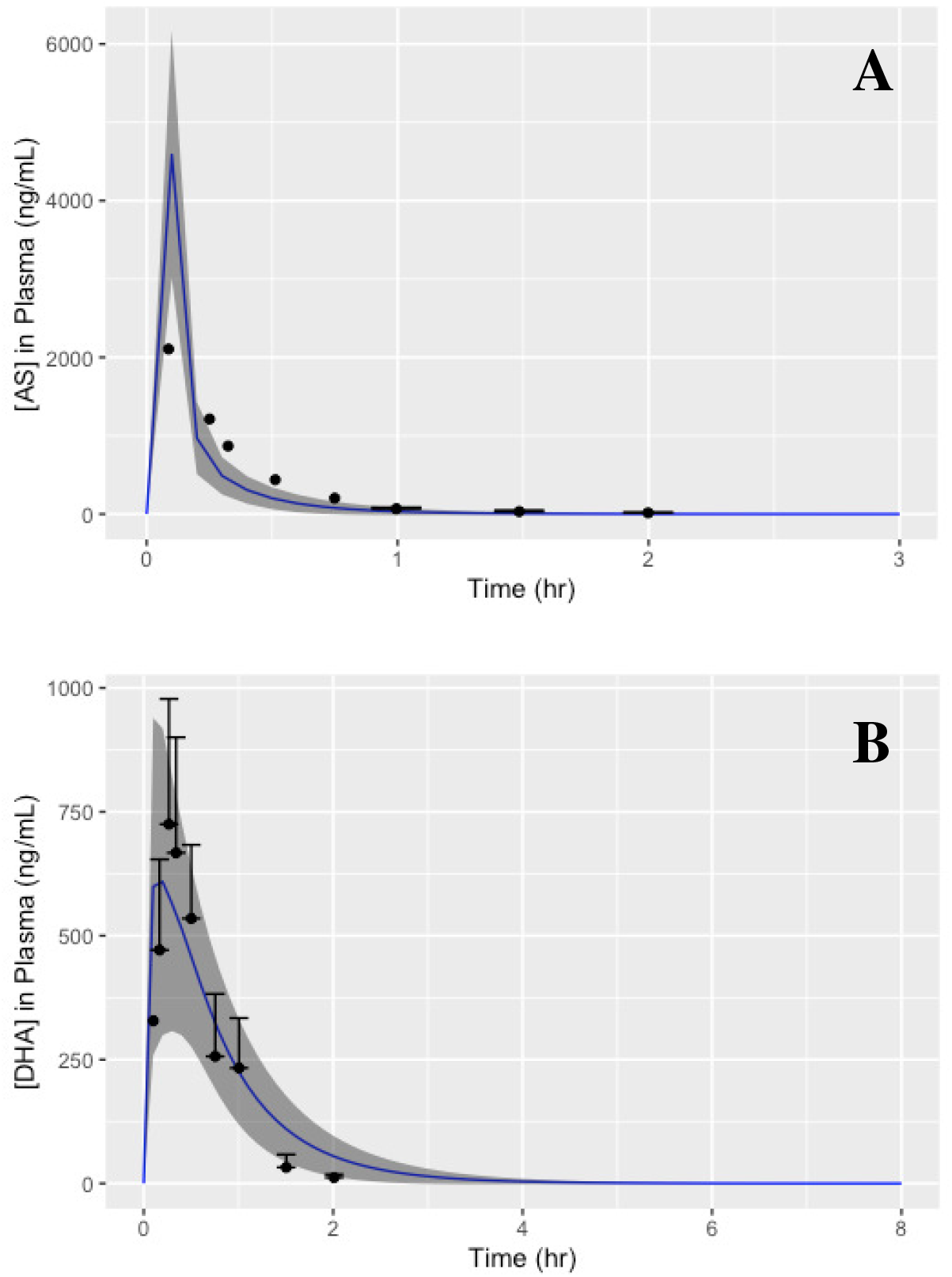
Model-predicted pharmacokinetics for unchanged AS (**A**) and unchanged DHA (**B**) in rat plasma following IV administration of AS at 10mg/kg. Simulations are co-plotted with data taken from the literature (**7**) for the purposes of model validation. Error bars were digitized from the sourced dataset.

**FIGURE 4.**
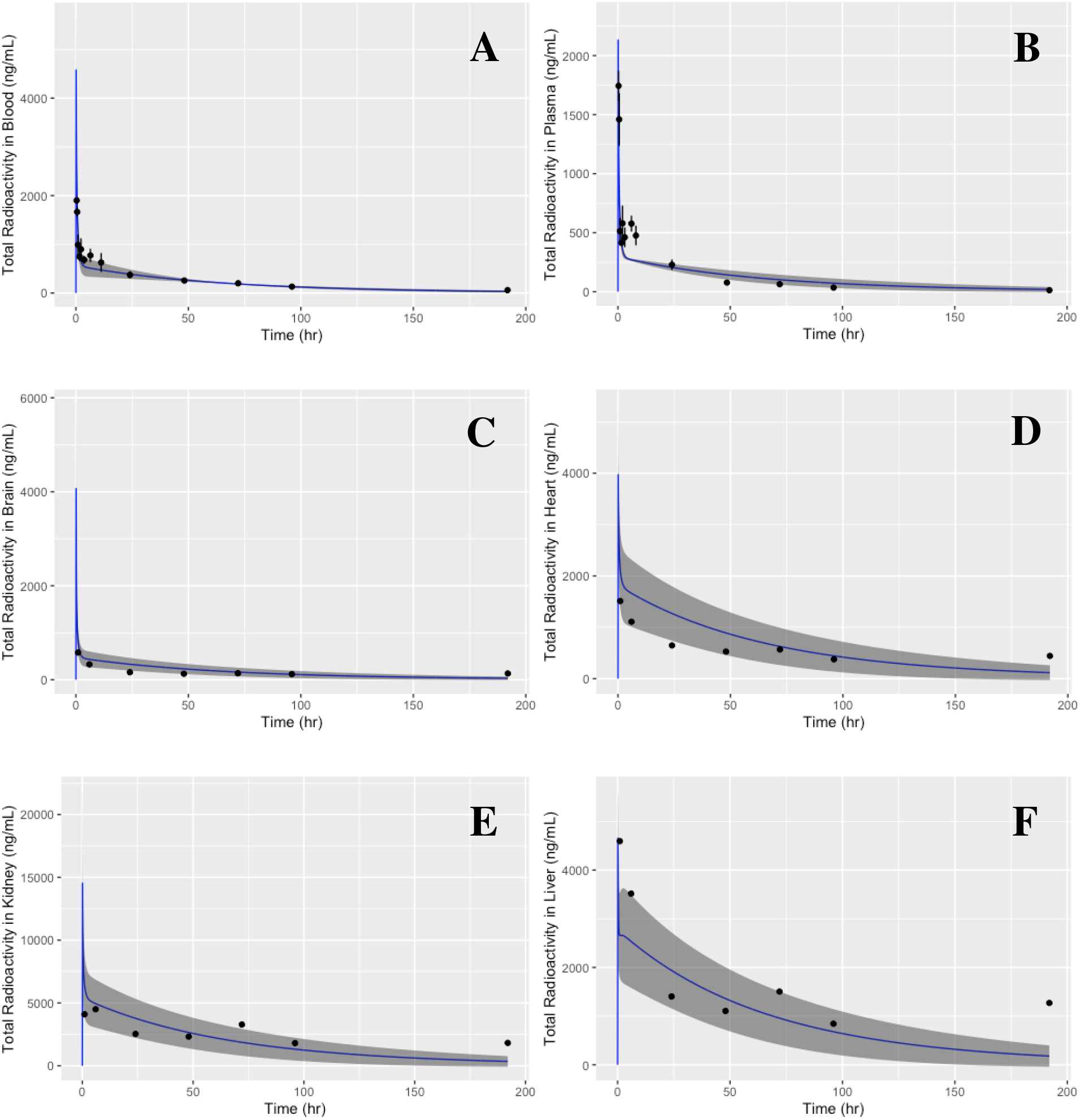
Model-predicted pharmacokinetics of TR concentrations in blood (**A**), plasma (**B**), brain (**C**), heart (**D**), kidney (**E**), and liver tissues (**F**) in rats following an intravenous dose of DHA at 3mg/kg. Simulations are co-plotted with data from the literature (**12**) for the purposes of model validation. Error bars for blood and plasma were digitized from the sourced dataset.

### Testing and validation of the H-PBPK model

Predictions from the H-PBPK model were assessed using the same method outlined above, through sampling the estimated human model parameters to produce a range of model outputs for comparison to the corresponding *in vivo* concentration-time data (**8**). The data selected for model comparison included AS and DHA concentrations in plasma resulting from IV administration of AS. This comparison is illustrated in **Figure 5**, which uses the same format as the plots described above. Further validation of the human model included simulations wherein AS was administered intravenously once every 24 hours for the span of 72 hours at 2mg/kg, 4mg/kg, and 8mg/kg. Concentration-time curves for plasma [DHA] were compared to the validation data (**11**), represented in **Figure 6**, which shows data (points) co-plotted with the mean PK profile of DHA (solid line) with 95% prediction intervals (greyed area) produced under this particular dosing scenario. IV doses of AS were simulated as a short bolus over the span of 2 minutes, and all validation data came from studies not used in the calibration step of the H-PBPK model.

**FIGURE 5.**
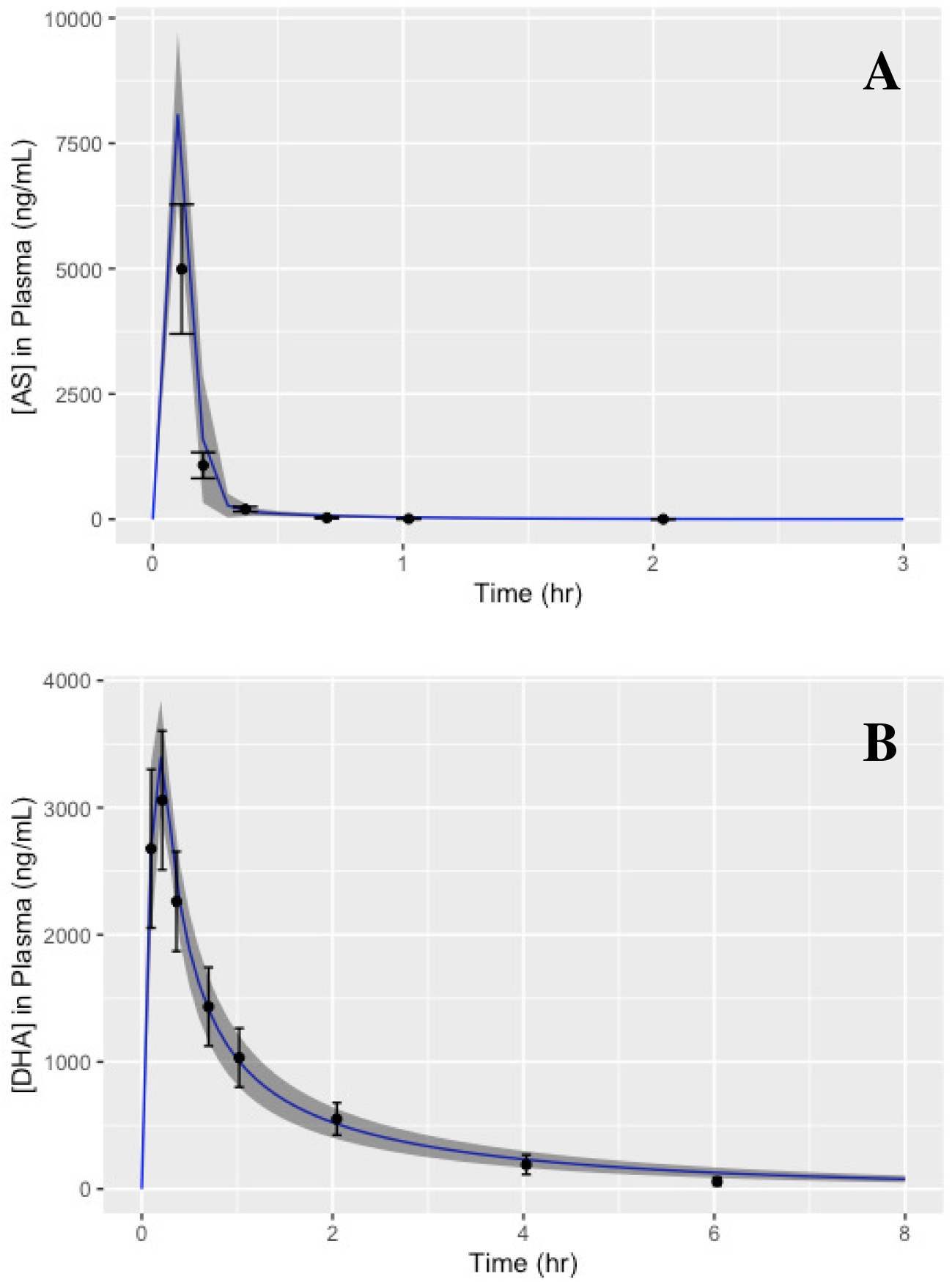
Model-predicted plasma pharmacokinetics of unchanged AS (**A**) and unchanged DHA (**B**) in patients with uncomplicated *Plasmodium falciparum* malaria following IV administration of AS at 2.4mg/kg. Simulations are co-plotted with data extracted from the literature (**8**) for model validation. Error bars were calculated from digitized points extracted from the sourced dataset.

**FIGURE 6.**
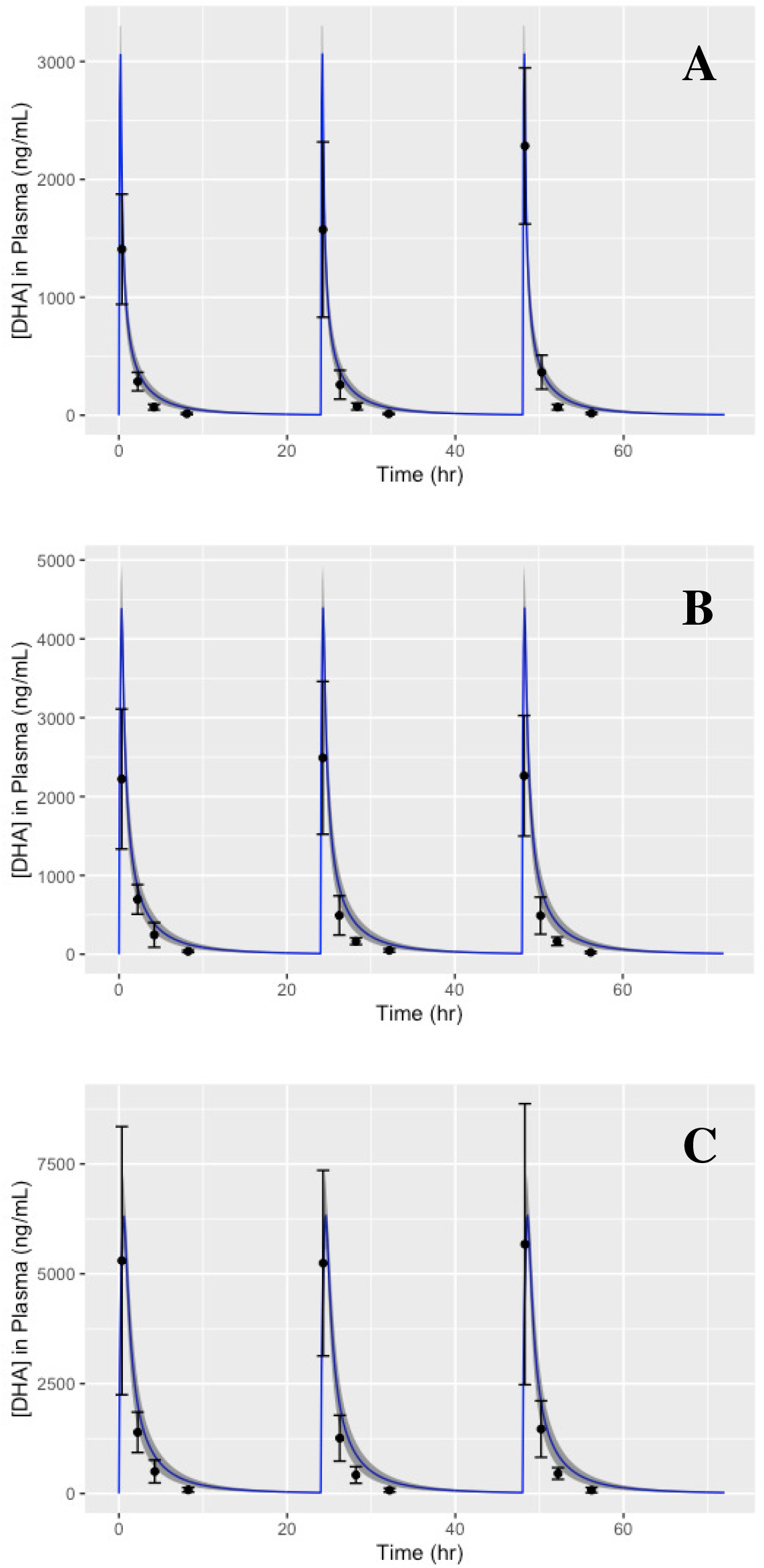
Simulations of the plasma pharmacokinetics of DHA in humans following a repeated dose schedule of IV_AS_ at 2mg/kg (**A**), 4mg/kg (**B**), and 8mg/kg (**C**) once every 24 hours for the span of 72 hours. Model-predictions are co-plotted with data pulled from the literature (**11**) for the purposes of model validation. Error bars were calculated from digitized points extracted from the sourced dataset.

As a final model characterization, PK parameters were calculated from a single simulation of the H-PBPK model utilizing the mean values for all estimated parameters calculated from their corresponding posterior distributions. PK parameters selected for model comparison were C_max_, half-life (t_1/2_), AUC, and mean residence time (MRT), as these measures were reported in the literature (**8**) and deemed important for consideration. This comparison can be seen in **Table 4**.

**TABLE 4.**
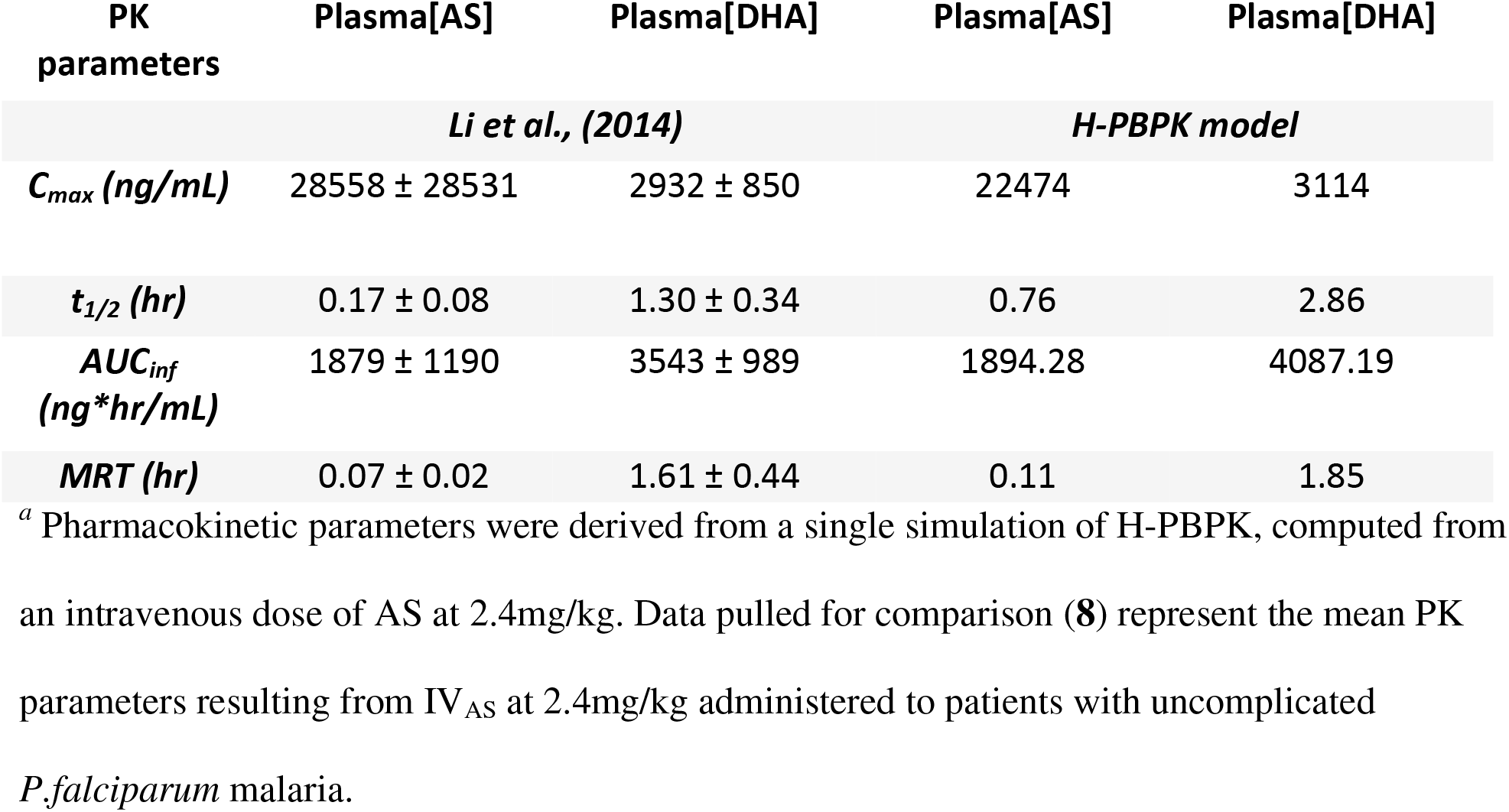
Computed pharmacokinetic parameters of AS and DHA for model comparison^*a*^

| PK parameters | Plasma[AS] | Plasma[DHA] | Plasma[AS] | Plasma[DHA] |
| --- | --- | --- | --- | --- |
|  | *Li et al., (2014)* |  | *H-PBPK model* |  |
| ***C<sub>max</sub>* (ng/mL)** | 28558 ± 28531 | 2932 ± 850 | 22474 | 3114 |
| ***t<sub>1/2</sub>* (hr)** | 0.17 ± 0.08 | 1.30 ± 0.34 | 0.76 | 2.86 |
| ***AUC<sub>inf</sub>* (ng\*hr/mL)** | 1879 ± 1190 | 3543 ± 989 | 1894.28 | 4087.19 |
| ***MRT* (hr)** | 0.07 ± 0.02 | 1.61 ± 0.44 | 0.11 | 1.85 |

<sup>a</sup> Pharmacokinetic parameters were derived from a single simulation of H-PBPK, computed from an intravenous dose of AS at 2.4mg/kg. Data pulled for comparison (8) represent the mean PK parameters resulting from IV<sub>AS</sub> at 2.4mg/kg administered to patients with uncomplicated *P.falciparum* malaria.

## Discussion

### Methodology

The PBPK models detailed herein utilized a system of physiological and biochemical descriptions, along with species mass balance equations, to make tissue-specific pharmacokinetic predictions for AS and its active metabolite, DHA, in relevant organs/tissues of both rats and humans. Data uncertainties and interstudy variability were incorporated in the results by estimating unknown parameter values within a hierarchical Bayesian framework. Simulations of both models were conducted using the posterior distributions resulting from model calibration and distributions of the mean value of other relevant parameters in order to quantify their effect on pharmacokinetic predictions.

### Testing and Verification

#### PBPK model for rats

Model predictions were generally in reasonable to very good agreement with the validation data. As shown in **Figures 3** and **4**, most of the data from multiple PK studies fell within the 95% prediction intervals produced by the model for blood, plasma, and tissue concentrations of both AS and DHA. Interestingly, the total radioactivity measures for blood and plasma species concentrations (**9, 10, 12**) indicated the presence of multiple concentration peaks, suggesting that these features are likely due, at least in part, to EHR of conjugated metabolites. This is consistent with findings from previous work on the metabolites of AS and DHA (**9, 19**). Though the model does include this biological mechanism, with the parameters estimated, the model does not recapitulate these PK features.

#### PBPK model for humans

In the process of H-PBPK model assessment, predictions were compared to single and repeated-dose data. **Figures 5** and **6** show that data from studies with various dosing regimens fell within the 95% prediction intervals produced by the H-PBPK model, demonstrating the model’s capacity to reasonably predict the plasma concentration-time profile of both AS and DHA. Likewise, as indicated in **Table 4**, the model-predicted PK parameters were found to fall within a single standard deviation of literature data, with the exception of t_1/2_ and MRT. Discrepancies between model predictions and PK experimental data could be attributed to simplified model assumptions on the metabolism of AS and DHA, and/or the use of parameters involved in AS metabolism taken from studies of other artemisinin derivatives (**26, 27**). Deficiencies in agreement between model predictions of t_1/2_ and MRT may also result from assumptions made about drug conjugation for both active compounds in the extrahepatic tissues listed previously (**24, 25**). Consideration of such processes would likely lead to an underprediction of t_1/2_ and an overprediction of MRT. Regarding convergence of the H-PBPK estimated parameters to a stationary distribution, the high R values pertaining to the PSRF of the posterior distributions of specific model parameters, namely *Km*_*AS3A4*_, *Km*_*AS3A5*_, *Cm*_*1*_, and *Cm*_*3*_ seem to indicate non-convergence. These results demonstrate a need for further refinement of the parameterization of the H-PBPK model, as described previously.

### Features and advantages of the present model

Unlike other PK models for AS and DHA (**6–12**), the present model provides information about tissue-specific drug concentrations and clearance characteristics. Predictions of drug levels close to the site of action are expected to aid investigators interested in both enhancing drug efficacy **(15, 48, 49**) and limiting the potential toxicity of artemisinin derivatives (**13**). A Though information about the dose response of artemisinins with respect to toxicity has not been established, it has been suggested that the risk lies in long-term availability rather than short-term peak concentrations (**13**). The current model addresses this concern by providing robust pharmacokinetic predictions for various key organs/tissues in the human body. Moreover, as with PBPK models in general, the present approach can facilitate a systematic examination of the anticipated pharmacokinetic effect of changes to dosing regimens and routes of administration. Finally, through the use of Bayesian inference, model parameters were estimated as distributions, allowing a quantitation of the effects of data and model uncertainly and intra-subject variability. With the listed advantages, the present model has the potential to aid in human dose optimization and help determine the extent to which pharmacokinetic endpoints depend on alterations to, and variability in, anatomical, physiological, and biochemical characteristics.

### Limitations of the present model

There are several limitations and deficiencies associated with the PBPK model described in this paper: (i) the present model does not recapitulate the presence of multiple concentration peaks that have been observed in experiments (**9, 10, 12**), though data uncertainty is relatively large in the datasets used; (ii) the model is not currently applicable to the analysis of drug combination therapies, which are are common; (iii) in the context of personalized medicine, as with almost all current PBPK models, the pharmacokinetic predictions contain too much uncertainty; and (iv) assumptions made about the metabolism of each active compound were based on *in vitro* data (**24, 25, 26**), which may not be reflective of *in vivo* metabolic characteristics.

### Future directions

Using the present model as a foundation, future work will be focused on adding additional anti-malaria agents (e.g., chloroquine, amodiaquine, mefloquine) to simulate combination therapies and quantify pharmacokinetic drug-drug interactions. Other enhancement will include integration of pharmacodynamic descriptions that encompass the growth and drug-induced killing kinetics of the malaria parasite, as well as descriptions of AS-induced toxicity in the relevant organs. Some of this work is already underway.

## Appendix

The equations listed in this section serve as the governing relationships for the PBPK model, which mathematically specify mass balance for each chemical species for each compartment, along with relevant biological phenomena involved in the disposition of AS and DHA. Blood flow rates to each tissue (Q_T_) were computed as Q_T_ = FQ_T_*Q_C_/Q_TC_, where Q_C_ symbolized cardiac output, scaled by Q_C_ = Q_CC_*BW^0.75^, FQ_T_ represented the fractional flow rate (fraction of cardiac output), and Q_TC_ represented the sum of all fractional flow rates. Organ/tissue volumes (V_T_) were scaled in a similar manner with V_T_ = FV_T_*BW/V_TC_, where FV_T_ symbolized the fractional volume of the given tissue, and V_TC_ represented the sum of all fractional tissue volumes. Furthermore, arterial blood concentrations entering each compartment denoted as C_ABu_ represent the fraction of drug unbounded to plasma proteins, while concentrations leaving each compartment in venous blood (C_TVB_) are calculated from tissue concentrations and tissue/plasma partition coefficients as follows:

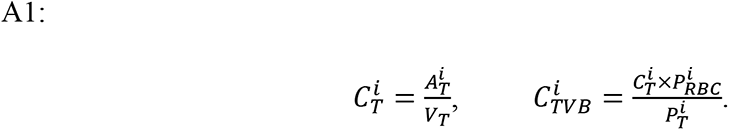

Here, the superscript “*i*” symbolizes any of the four the chemical species tracked by the model. 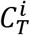 and 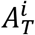 represent drug concentration and drug amount, respectively. 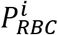 is the blood/plasma partition coefficient, and 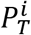 represents the partition coefficient of the given tissue. The differential equations for all non-eliminating organs, such as *brain* and *heart*, took the following form:

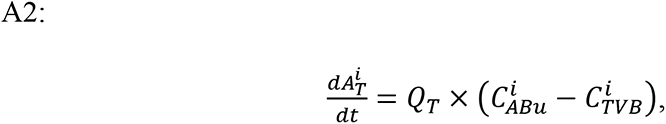

where the derivative 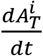 represents changes in the amount of drug with respect to time. All other compartments involved in the metabolism or excretion of drugs are described by the equations listed below. For reference, subscript “*j”* represents the various coefficients of metabolism, with *C*_*m*1_ pertaining to AS metabolism in venous blood, *C*_*m*2_ corresponding to AS conjugation in the liver and extrahepatic tissues, and *C*_*m*3_ being used to scale DHA conjugation in extrahepatic tissues.

### Muscle compartment

Equations for both active compounds (AS and DHA) in the muscle compartment took the following form:

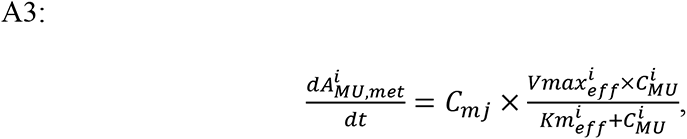

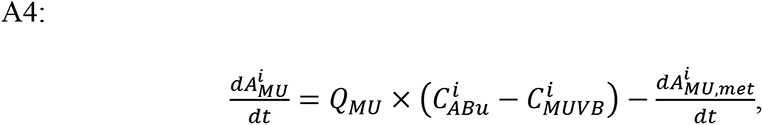

where the derivative 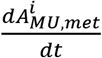 represented the amount of drug conjugated with respect to time, while the equations for both inactive compounds (AS-C and DHA-C) were computed with

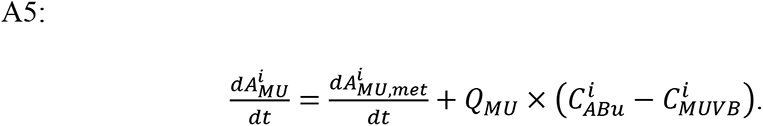

### Gut compartment

Equations for AS and DHA took the following form in the gut compartment:

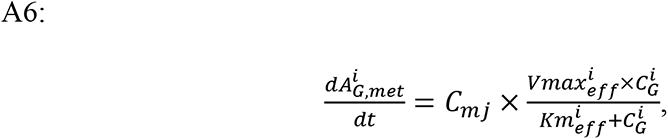

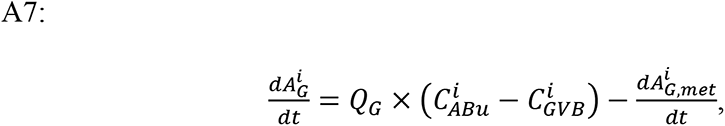

where 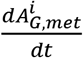 was used to compute the amount of drug conjugated with respect to time. The equations for AS-C and DHA-C were computed as follows:

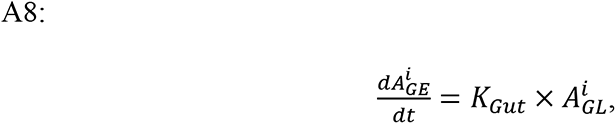

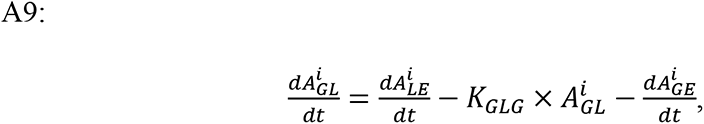

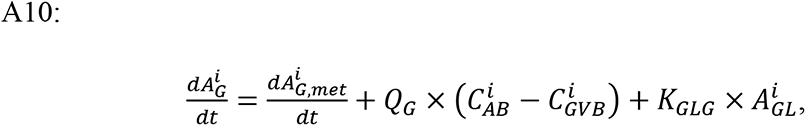

where 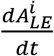 symbolized the amount of metabolite entering the intestinal lumen via biliary excretion. 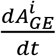 is the amount of drug being excreted from the gut lumen, 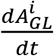 represents the rate of change in drug amount in the gut lumen, and 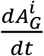 dictates the rate of change in drug amounts in the whole gut with respect to time.

### Kidney compartment

Equations for AS and DHA took the following form in the kidney compartment:

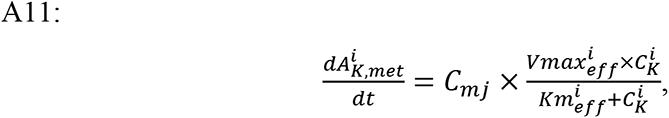

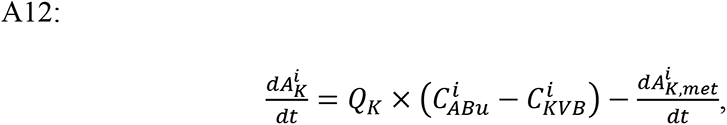

where 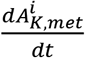 represents amount of drug metabolized with respect to time. Equations for AS-C and DHA-C were computed as follows:

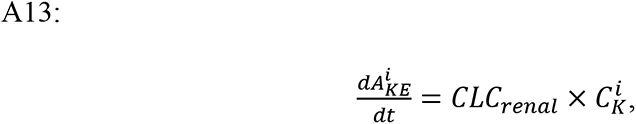

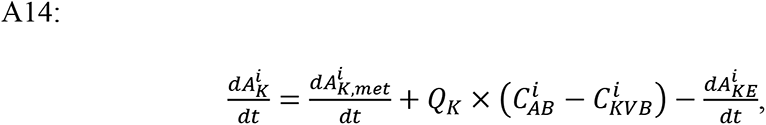

where 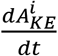 represented the amount excreted from the kidneys and 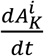 represented changes in the amount of drug with respect to time.

### Liver compartment

The equation for the conversion of AS to DHA in the liver was computed as the summation of all metabolic rates owing to the enzymes responsible for the metabolism of other artemisinin derivatives as follows:

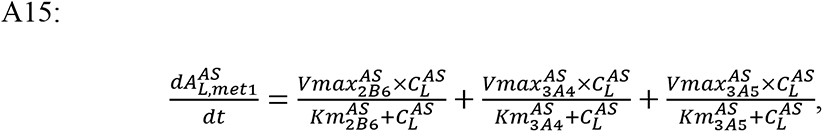

while conjugation of AS in the liver was computed with

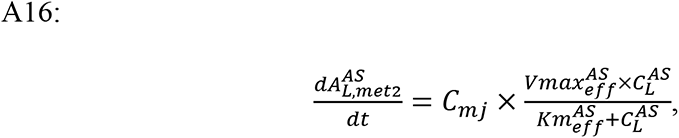

and changes in the amount of AS with respect to time were computed as

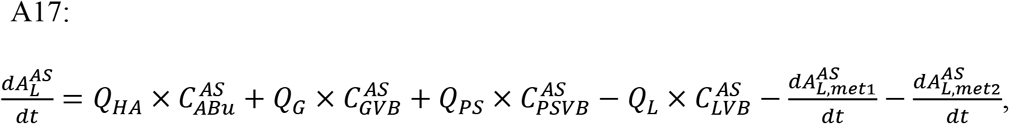

where *Q*_*G*_ + *Q*_*PS*_ is equivalent to the flow of blood from the portal vein. The conjugation of DHA in the liver compartment was represented by

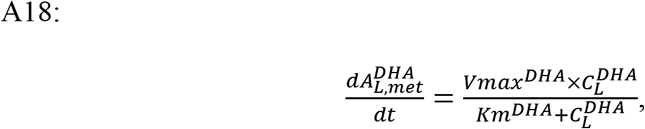

while changes in the amount of DHA with respect to time were computed as

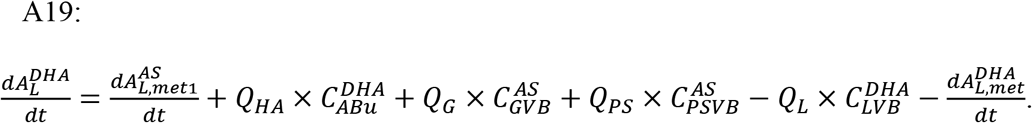

Biliary excretion of the conjugated metabolites was modeled by

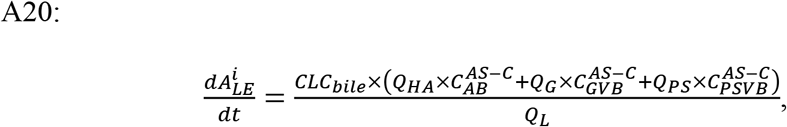

with changes in AS-C amounts computed as

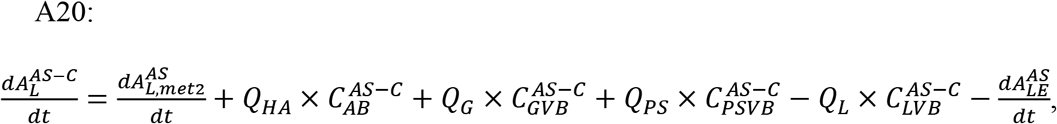

and changes in DHA-C amounts as

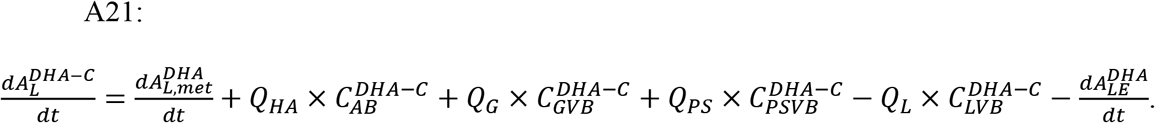

### Venous blood compartment

Changes in the amount of drug in the venous compartment were accounted for in the following equations:

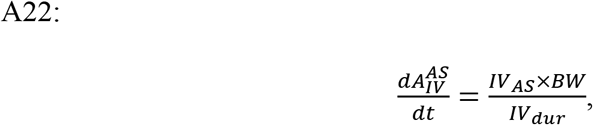

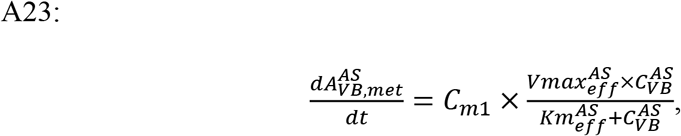

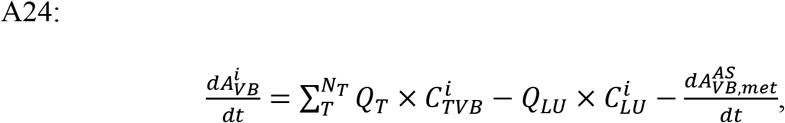

where 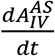 represented the IV dose calculation *IV*_*AS*_ symbolized the IV Dose (mg/kg), and *IV*_*dur*_ symbolized the duration of IV infusion. 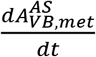 represented the metabolism of AS in venous blood and it was only applied to the appropriate equations. Calculation of total radioactivity (TR) concentrations in the various tissues was accomplished by summing the concentrations of every chemical compound in the given tissue. This computation was carried out for the span of each model simulation as

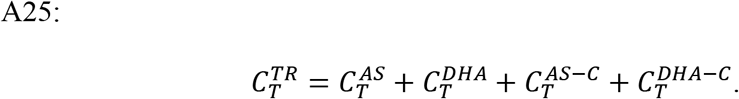

All drug amounts were computed in units of mg, while drug concentrations were computed in units of mg/L. Model outputs used for comparison to the validation dataset were converted to the necessary units before being co-plotted with the data.

